# mAML: an automated machine learning pipeline with a microbiome repository for human disease classification

**DOI:** 10.1101/2020.02.11.943316

**Authors:** Fenglong Yang, Quan Zou

**Affiliations:** Institute of Fundamental and Frontier Sciences, University of Electronic Science and Technology of China, Chengdu, China

## Abstract

Due to the concerted efforts to utilize the microbial features to improve disease prediction capabilities, automated machine learning (AutoML) systems designed to get rid of the tediousness in manually performing ML tasks are in great demand. Here we developed mAML, an ML model-building pipeline, which can automatically and rapidly generate optimized and interpretable models for personalized microbial classification tasks in a reproducible way. The pipeline is deployed on a web-based platform and the server is user-friendly, flexible, and has been designed to be scalable according to the specific requirements. This pipeline exhibits high performance for 13 benchmark datasets including both binary and multi-class classification tasks. In addition, to facilitate the application of mAML and expand the human disease-related microbiome learning repository, we developed GMrepo ML repository (GMrepo Microbiome Learning repository) from the GMrepo database. The repository involves 120 microbial classification tasks for 85 human-disease phenotypes referring to 12,429 metagenomic samples and 38,643 amplicon samples. The mAML pipeline and the GMrepo ML repository are expected to be important resources for researches in microbiology and algorithm developments.

**Database URL:** http://39.100.246.211:8050/Home

## 1 Introduction

Machine learning (ML) models are crucial for data-driven medical research and translation, such as microbiome-based disease diagnosis or prognosis (1–3). As domain classification tasks are often context-dependent, no single data preprocessing method and ML strategy can handle all prediction issues. Due to the tedious nature of customizing ML tasks by domain scientists, several well-known automated machine learning (AutoML) systems, including Auto-WEKA (4), Auto-sklearn (5), and Auto-Net (6) have emerged to address the famous CASH (automatically and simultaneously choosing a ML algorithm and setting its hyper-parameters to optimize empirical performance) problem (7). While autoML systems are mature enough to enhance the prediction accuracy, few of them are designed to meet the specific requirements of microbial classification tasks (8). Focusing on the scenario of microbiome-associated phenotype prediction and considering the benefit of data preprocessing methods in regard to ML estimators (1), an autoML pipeline named mAML was developed here to automatically generate an optimized ML model that exhibits sufficient performance for a personalized microbial classification task.

The mAML pipeline possesses several advantages. Specifically, i) The mAML pipeline can efficiently and automatically build an optimized, interpretable and robust model for a classification task. ii) The mAML pipeline is deployed on a web-based platform (the mAML web server) that is user-friendly, flexible and has been designed to be scalable according to user requirements. iii) The pipeline can be applied to both binary and multi-class classification tasks. iv) The pipeline is data-driven and it can be easily extended to the multi-omics data or other data types if only the domain-specific feature matrix is provided.

Furthermore, we developed a microbiome learning repository from the GMrepo database (9). GMrepo (data repository for Gut Microbiota) is a database of curated and consistently annotated human gut metagenomic data, which contains 58903 human gut samples/runs, including 17618 metagenomes and 41285 amplicons from 253 projects concerning 92 phenotypes. GMrepo consistently processed and annotated the collected samples and manually curated all possible related meta-data of each sample/run. It organized the samples according to their associated phenotypes and offered the taxonomic (genus and species level) abundance information for all samples of high quality. Due to the necessities of aggregating samples across studies and appropriately handling candidate confounders in curating classification tasks, it’s reasonable to develop a machine learning repository dedicated to human-disease associated microbial classification tasks from GMrepo. Hence, we present the GMrepo ML repository (GMrepo Microbiome Learning repository), a public repository of 120 microbial classification tasks developed from the GMrepo database, which involves 38,643 amplicon samples referring to 71 disease phenotypes and 12,429 metagenomic samples covering 49 disease phenotypes. The files in the GMrepo ML repository can be downloaded and directly submitted to the mAML server or they can be imported into the phyloseq pipeline (15) for rapid, reproducible and interactive exploration of microbiome data.

The source code and benchmark datasets for the mAML pipeline are available at https://github.com/yangfenglong/mAML1.0. The docker image of the pipeline can be pulled from https://hub.docker.com/r/yangfenglong/dash_webserver. The GMrepo ML repository is freely available at http://39.100.246.211:8050/Dataset. The source code for the construction of the repository is available at https://github.com/yangfenglong/mAML1.0/blob/master/datasets/GMrepo_datasets/GMrepo.ipynb.

## 2 Method

The mAML pipeline is developed completely in Python, and the workflow is represented in Figure 1. First, the features that exhibit a prevalence lower than the threshold of 20% by default in all classes were filtered, as low prevalence features are usually not promising for use in the analysis of gut microbiota that possesses thousands of features across samples. Second, the irrelevant and redundant features were further removed using the mRMR (Minimum redundancy Maximum relevance algorithm) method (10) while retaining the leading features (top 50 features by default) to allow for the improvement and comprehensibility of the model and to save computer resources. Then, the class imbalance problem was compensated by using SMOTE (Synthetic Minority Over-sampling Technique) (11), as imbalanced datasets containing overrepresented or underrepresented data can induce bias in prediction. Finally, the pipeline will automatically determine the optimized hyper-parameters for all classifiers, including the best preprocessors for non-tree-based classifiers, using parallel grid searches. The use of appropriate data preprocessing methods such as feature scaling is important when used in combination with normality assumed algorithms such as metric-based, gradient-based, and distance-based estimators, and it has no influence on tree-based models (1). Hence, the optimized hyper-parameters for those non-tree based models were explored while considering data preprocessing methods as one of the important hyper-parameters. Nested cross-validation (12) was used to avoid overfitting for each model in regard to training data, and 11 candidate scoring metrics, with accuracy as the default, were applied for model evaluation. In total, 10 data preprocessing methods and 13 classifiers, primarily derived from python machine learning package scikit-learn (1), were involved. Considering the interpretation requirements for the subsequent microbial studies, only those white-box preprocessors and classifiers were incorporated.

**Figure 1.**
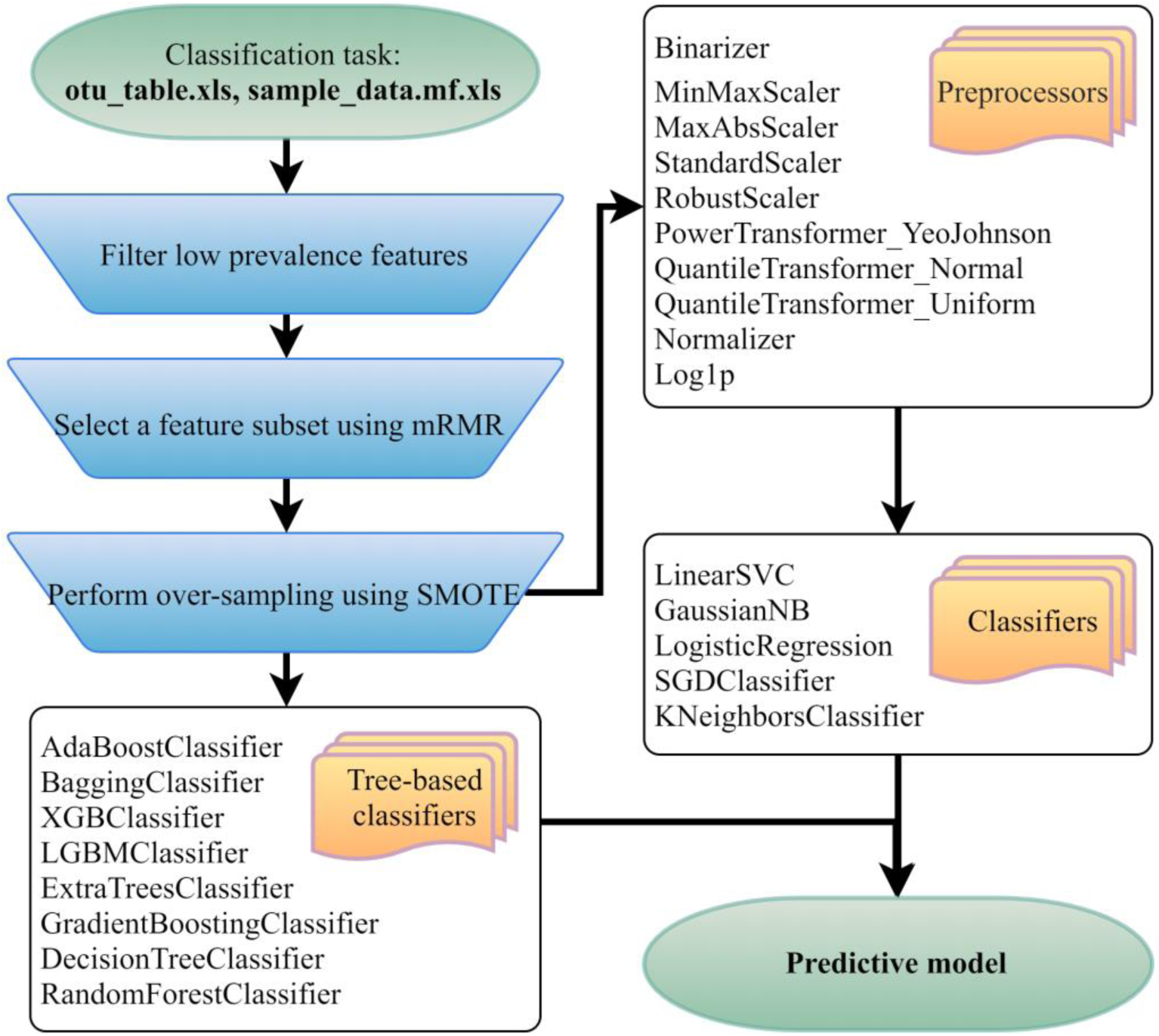
Flowchart of the mAML pipeline. Two files indicated at the beginning of the pipeline should be provided. Operation steps before training are indicated in the blue inverse-trapezoids.

## 3 Utility of mAML

### Overview

To facilitate the use of the mAML pipeline, we developed the mAML web server, which is a web-based machine learning system that can generate models in a user-friendly, flexible, and scalable way. The main points regarding the implementation of the server are described as follows and the details can be accessed on the server “Help” page.

### Web Server Implementation

Here we will introduce the key points to navigate the server.

### Submit a task

A typical classification task can be submitted to the mAML web server by the following steps (Figure 2). First, users can choose the example datasets or upload datasets of their own to start the pipeline. Second, the input features will be filtered at the specified threshold (by default, taxonomic features with a percentage lower than 20% in all classes were disregarded in this work). Third, the most relevant features will be selected using mRMR (top 50 features by default), and this option can be deselected if there’s no need to downsize the number of features. Fourth, the unbalanced datasets will be rebalanced using SMOTE, and this option can be turned off if it’s unnecessary to perform SMOTE. Finally, the default parameters and hyperparameters of the preprocessors and classifiers could be reconfigured, and the adding and pruning of any preprocessor or classifier is supported. The grid search settings for the hyperparameters can be altered in the dict function of each classifier. By default, the pipeline will search the best combination of preprocessors and non-tree based classifiers and simultaneously optimize the hyperparameters for all classifiers. The parameters of nested cross-validation, the metrics for model evaluation and the number of parallel processes are also tunable.

**Figure 2.**
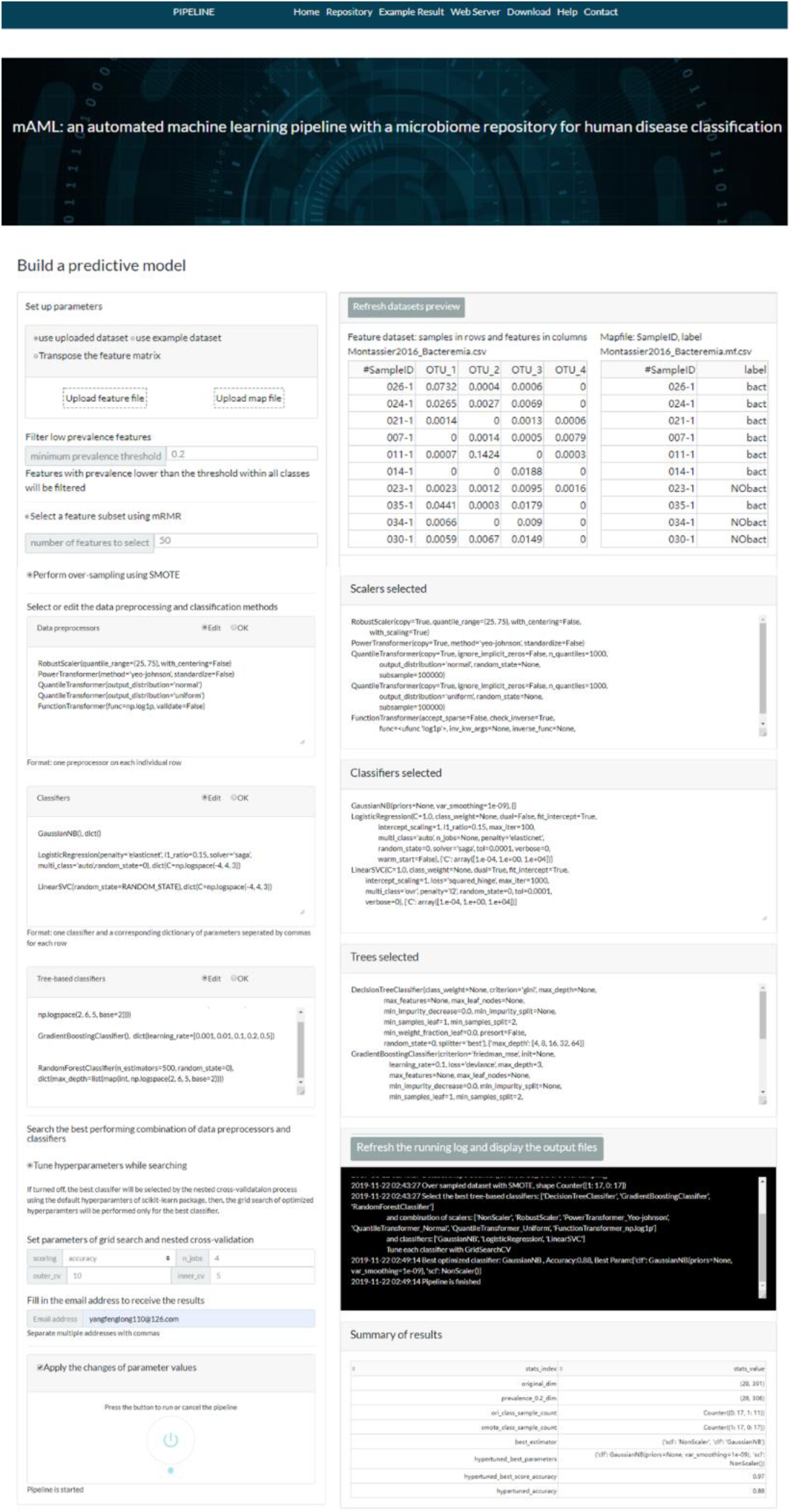
The task submission page of the mAML server. The settings for each step can be reconfigured and the status of the run can be checked by pressing the refresh button.

When the email address is filled in and all the above settings are confirmed, the user can start the pipeline and it will run in the background. The running process can be checked by pressing the refresh button on the server page.

### Preview the result files and download

Once the run is completed, the compressed result will be automatically sent to the predefined e-mail address or can be downloaded from the “Web Server” page of the server. The user can preview the interaction diagram (Figure 3) for the performance of all the candidate models on the server. For each task, the pipeline automatically outputs the visualization results for the optimal model, including the heatmap of confusing matrix (Figure 4A), the classification report (Figure 4B), ROC curve (Figure 4C) and the histogram of top features (Figure 4D, default: 20), which can be investigated in further study.

**Figure 3.**
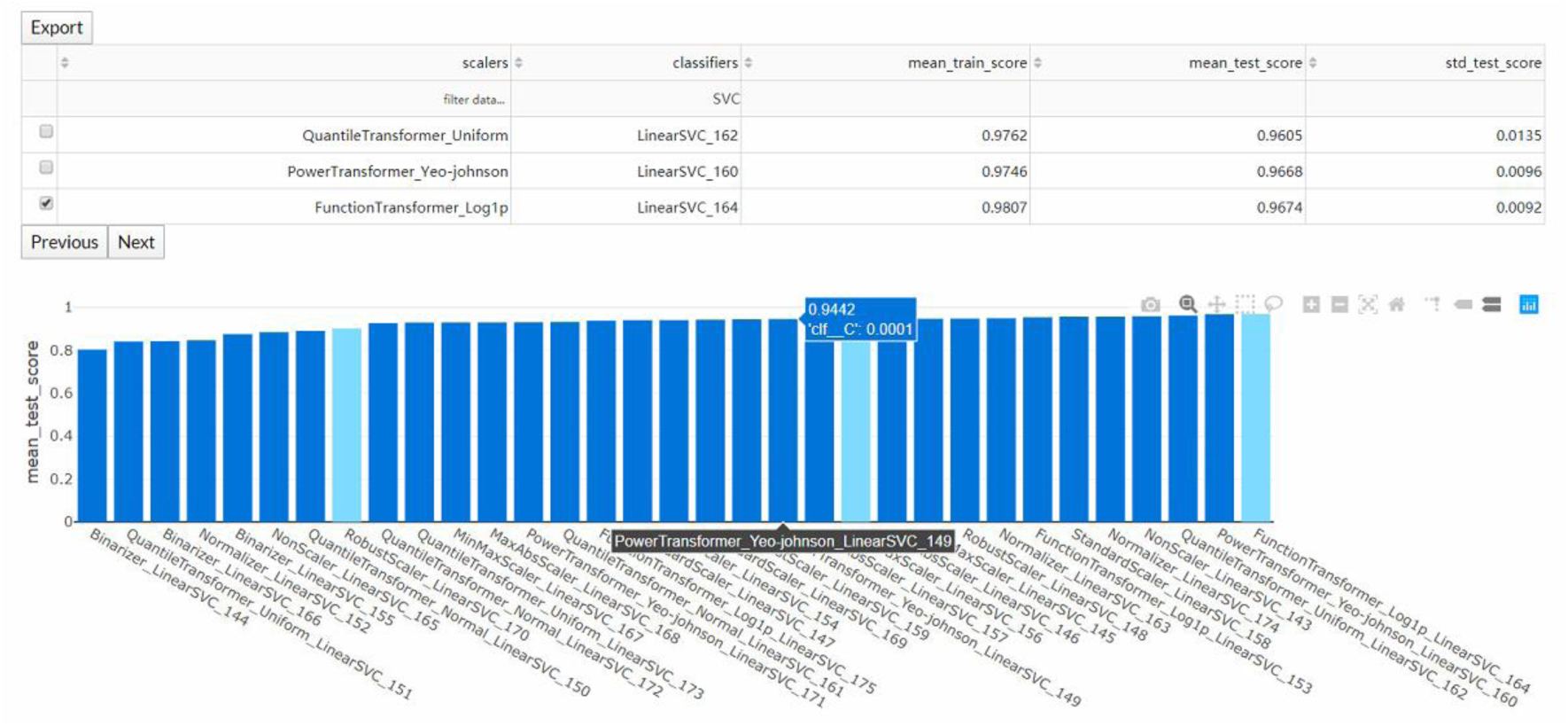
The interaction diagram for the performance of all the candidate models. The users can screen the candidate models by the scalers, classifiers, mean training score, mean test score, and standard test score.

**Figure 4.**
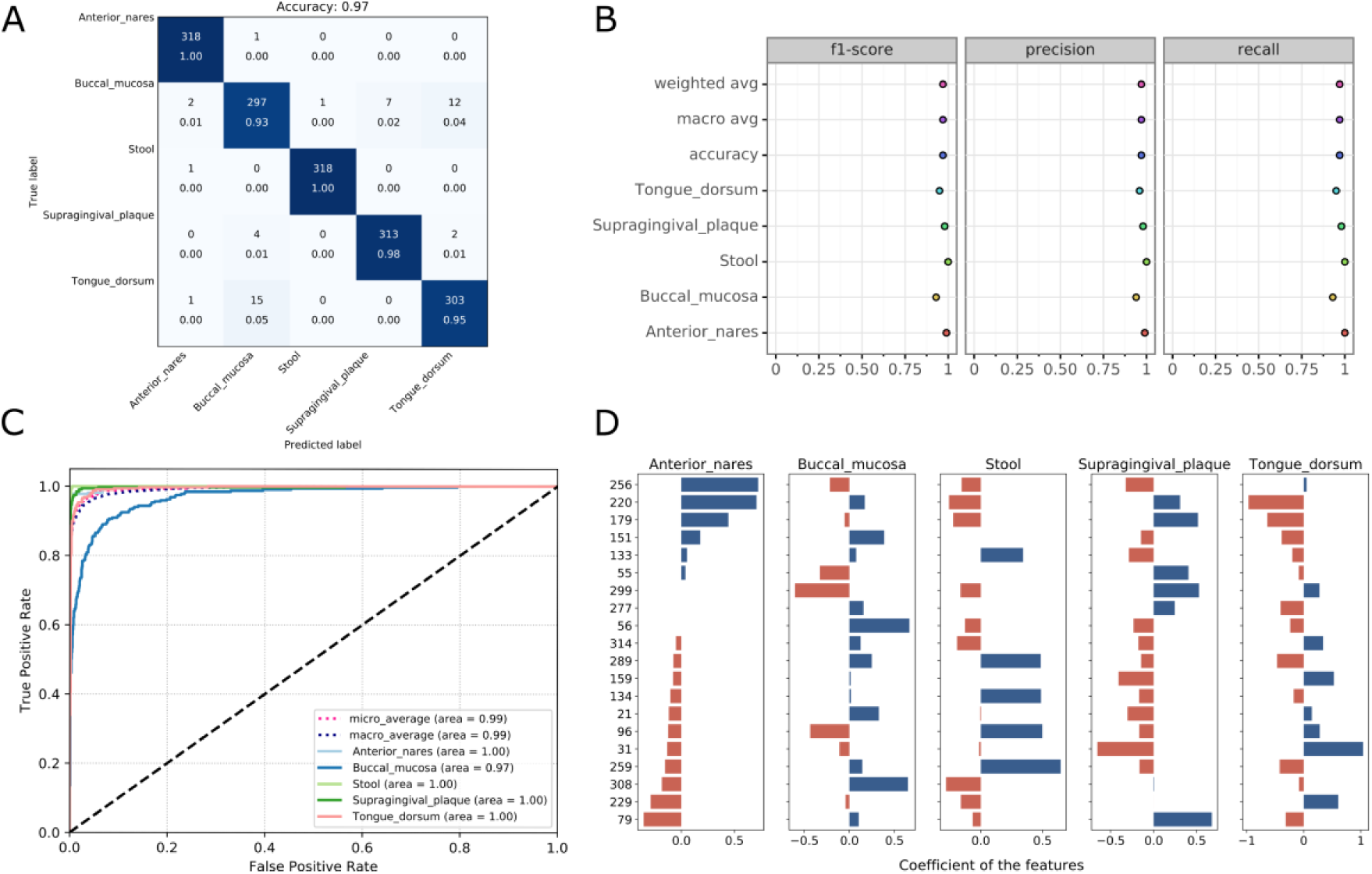
Visualizations for the optimized model: the heatmap for confusing matrix (Figure 4A), the classification report (Figure 4B), ROC curve (Figure 4C) and the histogram for top features (Figure 4D, default: 20). Note that, in the case of tree-based models, the feature importance will be provided instead of the feature’s coefficient in the histogram.

The result is reproducible within a container started based on the docker image or via the webserver. An example result is represented on the “Example Result” page of the server.

### Make New predictions

The user can feed new data to the existing model or upload a previously trained model to get new predictions (Figure 5).

**Figure 5.**
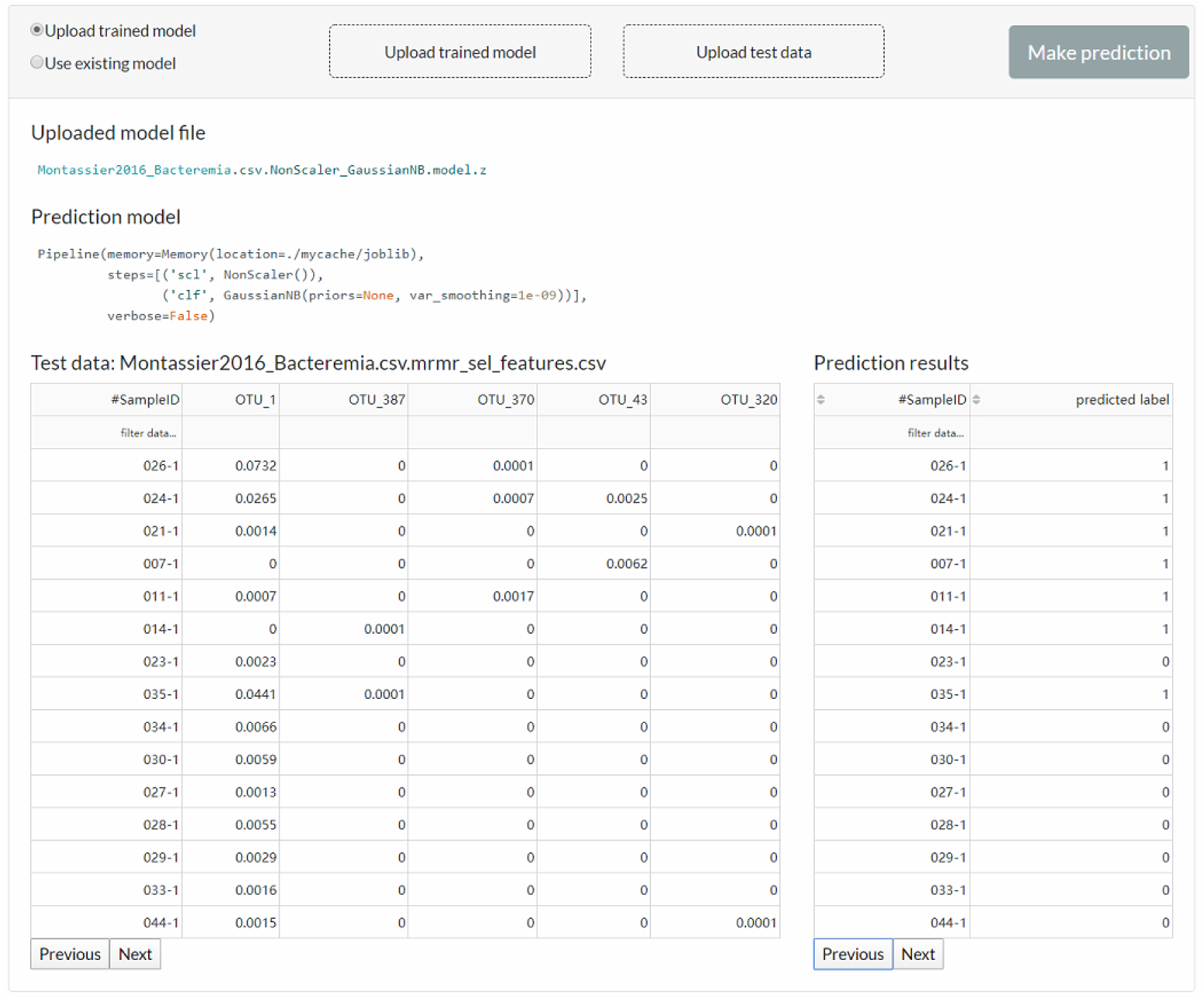
The web server interface for the users to reuse the existing model or upload a previously trained model to make predictions.

### Performance of mAML

The performance of the pipeline was investigated by performing analyses on 13 human microbiome datasets that are publicly available and appropriate for benchmarking, which involve 7 binary classification tasks (13) and 6 multi-class classification tasks (14). These datasets, including 11 amplicon datasets and 2 metagenomic datasets, vary across microbiome-related phenotypes and cover 2,471 samples, which can be retrieved from the “Repository” page of the server or GitHub repository.

The models proposed by mAML outperform most of the models in the original studies (Table S1, Figure 6), confirming the robustness and reliability of this method. Since only the “white box” preprocessors and classifiers are involved in the prediction, the optimized model selected for each task is interpretable and the top features indicated by the model merit further study. The detailed results for the candidate models and optimized model of each task are available at the GitHub repository.

**Figure 6.**
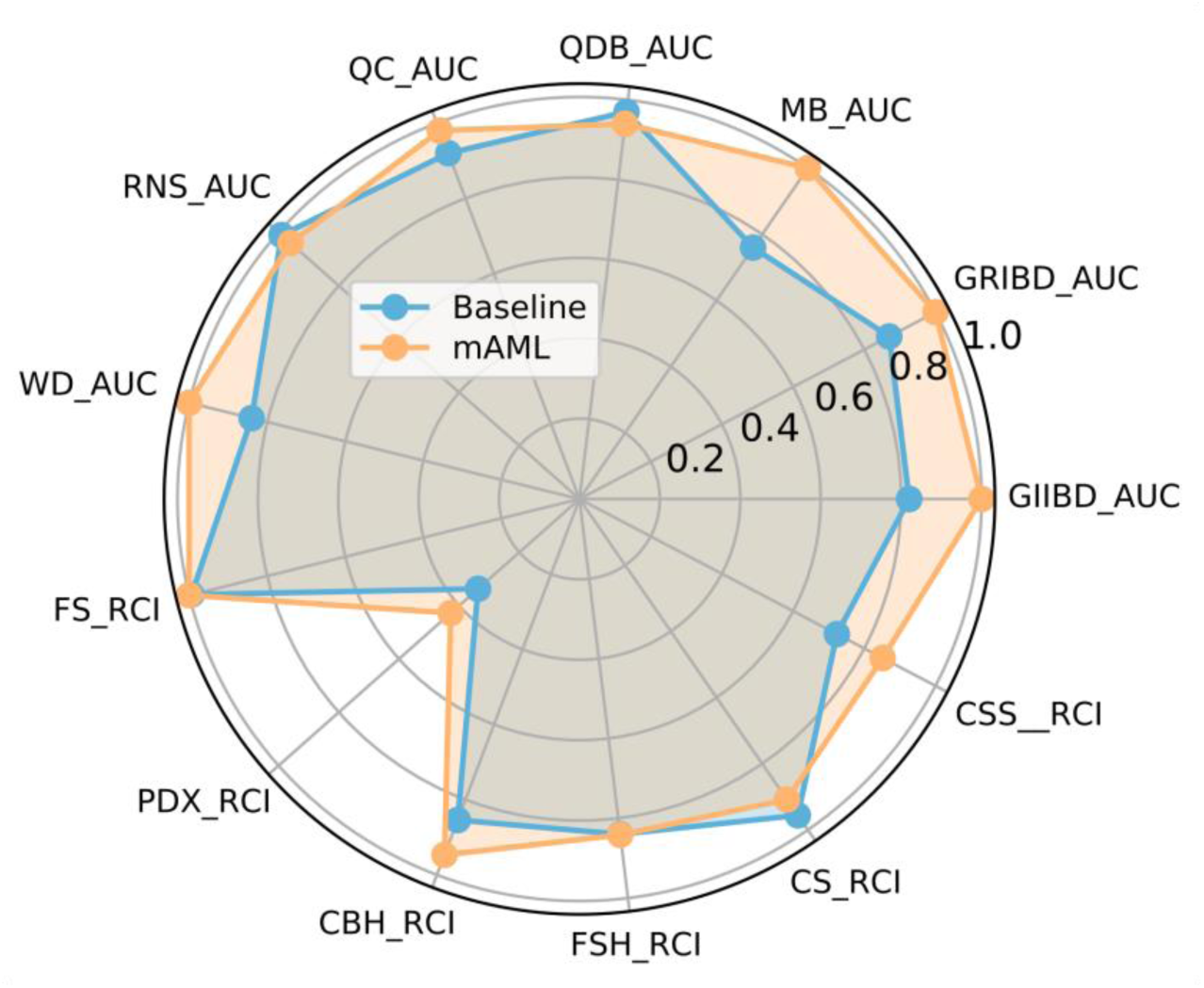
The performance comparison of the mAML proposed models against the baseline. The labels are connected abbreviations with an underline between the name of the database and the metric used in the original study.

### GMrepo ML repository

Furthermore, we developed a GMrepo ML repository (GMrepo Microbiome Learning repository) from the GMrepo database to facilitate the utilization of mAML and expand the microbiome learning repository related to human disease. The framework of the GMrepo ML repository construction is presented in Figure 7 and the details are described as follows.

**Figure 7.**
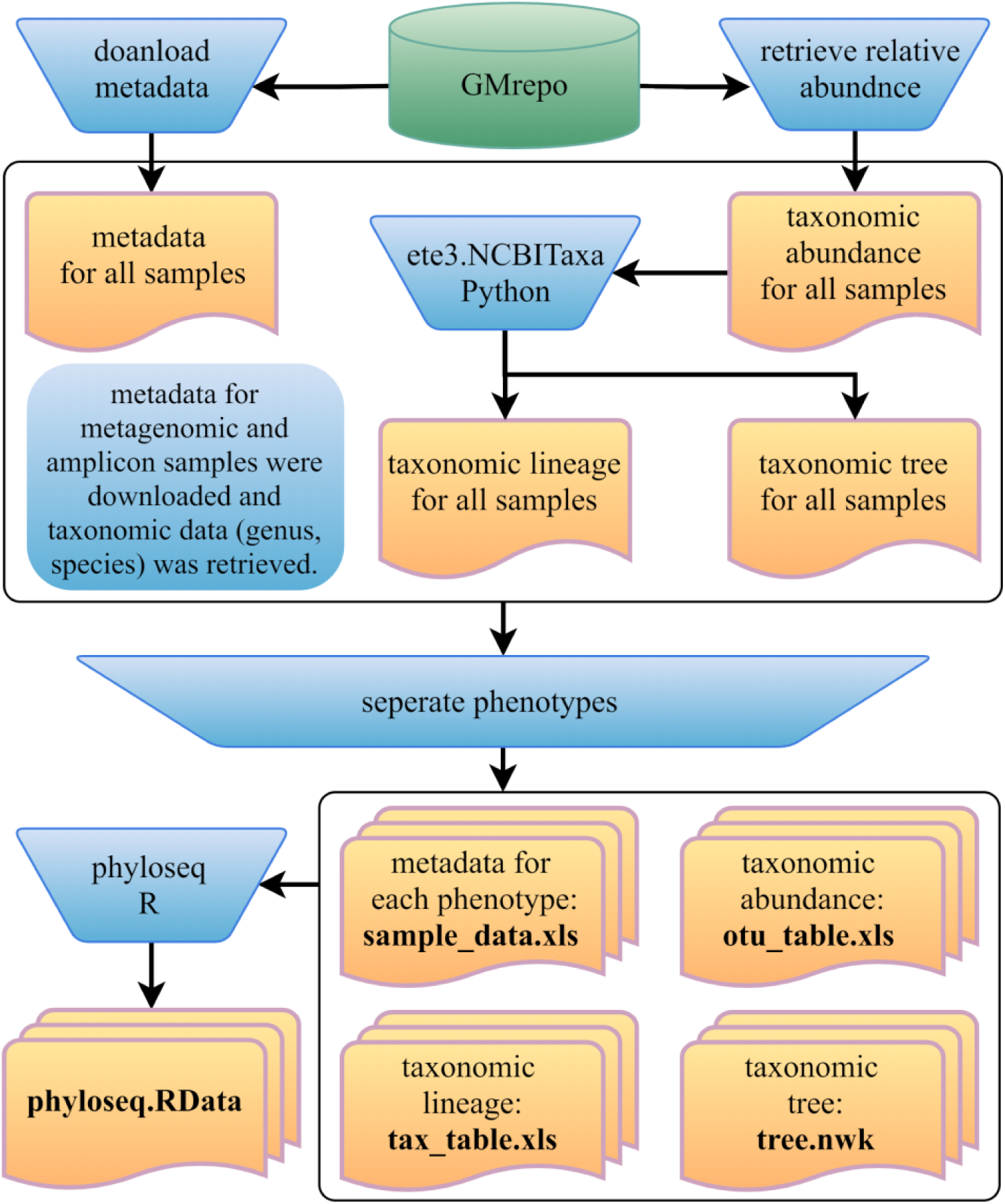
Framework of the GMrepo ML repository construction. Operation steps are indicated in the blue inverse-trapezoids. Files with names in bold are all contained in the repository, and they can be retrieved from the “Repository” page of the server or from the GitHub.

We downloaded the metadata for all samples that passed the QC procedure (QCStatus = 1) from the GMrepo website and retrieved the taxonomic abundance information (including the genus and species level) for all the metagenomic and amplicon samples respectively using the RESTful APIs of GMrepo. Taxa with the scientific name labeled as unknown or Others were deleted from the taxonomic abundance table since they are not meaningful features. Taxonomic lineage and tree were retrieved according to the NCBI taxonomy id of each taxon by using the ETE3 python module. Then four files (taxonomic abundance table, metadata table, taxonomic lineage table and taxonomic tree file) were obtained for all the metagenomic and amplicon samples respectively, as represented in Figure 7. Each file was then divided into subfiles according to different phenotypes and only those phenotypes with healthy control samples are preserved. Totally, the repository involves 12,429 metagenomic samples covering 49 disease phenotypes and 38,643 amplicon samples referring to 71 disease phenotypes. For each phenotype, the taxonomic abundance table and metadata table can be directly submitted to the mAML server to build an optimized model for disease prediction. Additionally, by using the phyloseq R package (15), all the subfiles files from each phenotype can be taken as components to build the phyloseq-class object (phyloseq.RData), which can be imported into the Shiny-phyloseq web application (16) for subsequent interactive exploration of microbiome data.

The users can apply the mAML pipeline to their interested datasets in the GMrepo ML repository. The datasets can also be merged with their own samples to perform meta-analysis. Moreover, multiple feature types such as metabolites and metatranscripts are encouraged to be integrated with the taxonomic or functional features from metagenomics to build a multi-omics-feature based model to the target disease.

## 4 Conclusion

Considering the tedious work and context-dependent nature of manually performing the microbial classification tasks, we developed an autoML pipeline, namely mAML, which can rapidly and automatically generate an optimized and interpretable model with sufficient performance for binary or multi-class classification tasks in a reproducible way. The pipeline is deployed on a web-based platform and the mAML server is user-friendly, flexible, and has been designed to be scalable according to user requirements.

We highlight the reliability and robustness of mAML with its high performance on 13 benchmark datasets. Being data-driven, the mAML pipeline can be easily extended to the multi-omics data of microbes and other data types if only the domain-specific feature data are supplied. Moreover, we constructed the GMrepo ML repository of 120 microbial classification tasks for 85 disease phenotypes, which facilitates the application of mAML and is expected to be an important resource for algorithm developers.

## Supplementary data

Supplementary data are available at *Database* Online.

