## Supplementary material for "mAML: an automated machine learning pipeline with a microbiome repository for human disease classification": Table S1

Supplementary Table 1 Details of the benchmark datasets and the performance comparison between the models proposed by mAML pipeline and the models of the original studies.

| Datasets with abbreviations in brackets | # of samples | # of features | # of classes | Labels in benchmark | **Baseline _models** | **mAML_ models** | Classification tasks with number of samples in brackets for each class |
|---|---|---|---|---|---|---|---|
| Gevers et al. (2014) Ileum IBD (GIIBD) | 140 | 943 | 2 | GIIBD_AUC | **0.82** | **1.00** | IBD: no (62), CD (78) |
| Gevers et al. (2014) rectum IBD (GRIBD) | 160 | 943 | 2 | GRIBD_AUC | **0.87** | **1.00** | IBD: no (92), CD (67) |
| Montassier et al. (2016) Bacteremia (MB) | 28 | 541 | 2 | MB_AUC | **0.76** | **1.00** | Bacteremia: bact (17), NObact (11) |
| Qin et al. (2012) Diabetes (QDB) | 124 | 11,880 | 2 | QDB_AUC | **0.97** | **0.94** | Diabetes: Y (65), N (59) |
| Qin et al. (2014) Cirrhosis (QC) | 130 | 8,483 | 2 | QC_AUC | **0.92** | **0.98** | Cirrhosis: Cirrhosis (68), Healthy (62) |
| Ravel et al. (2011) Nugent score (RNS) | 342 | 586 | 2 | RNS_AUC | **0.99** | **0.96** | Nugent_score_category: low (245), high (97) |
| Wu et al. (2011) Diet (WD) | 96 | 292 | 2 | WD_AUC | **0.84** | **1.00** | Diet: HighFat (40), LowFat (45) |
| Fierer et al. (2010) Subject (FS) | 104 | 294 | 3 | FS_RCI | **0.99** | **1.00** | 3 subjects: (40, 33, 31) |
| Yang et al. (2010) Diagnosis (PDX) | 200 | 5955 | 4 | PDX_RCI | **0.34** | **0.43** | Esophagitis: normal (28), reflux esophagitis (36), Barrett's esophagus (84), esophageal adenocarcinoma (52) |
| Costello et al. (2009) Body Habitat (CBH) | 552 | 1454 | 6 | CBH_RCI | **0.85** | **0.95** | Body habitats: skin (357), oral cavity (46), External Auditory Canal (44), Hair (14), Nostril (46), Feces (45) |
| Fierer et al. (2010) Subject x Hand (FSH) | 98 | 294 | 6 | FSH_RCI | **0.84** | **0.84** | Subject and left/right hand: (20, 18, 17, 14, 16, 13) |
| Costello et al. (2009) Subject (CS) | 140 | 464 | 7 | CS_RCI | **0.96** | **0.91** | 7 subjects: (20, 20, 20, 20, 20, 20, 20) |
| Costello et al. (2009) Skin Sites (CSS) | 357 | 600 | 12 | CSS__RCI | **0.72** | **0.85** | Skin sites: external nose (14), forehead (32), glans penis (8), labia minora (6), axilla (28), pinna (27), palm (64), palmar index finger (28), plantar foot (64), popliteal fossa (46), velar forearm (28), umbilicus (12) |

AUC: area under thecurve; RCI: relative classifier information; IBD: inflammatory bowel disease; CD: Crohn’s disease
